# Bclaf1 critically regulates the type I interferon response and is degraded by alphaherpesvirus US3

**DOI:** 10.1101/392555

**Authors:** Chao Qin, Rui Zhang, Yue Lang, Anwen Shao, Aotian Xu, Wenhai Feng, Jun Han, Mengdong Wang, Wanwei He, Cuilian Yu, Jun Tang

**Affiliations:** State Key Laboratory of Agrobiotechnology and College of Veterinary Medicine, China Agricultural University, Beijing 100193, China; Department of Microbiology and Immunology, College of Biological Sciences, China Agricultural University, Beijing 100193, China

## Abstract

Type I interferon response plays a prominent role against viral infection, which is frequently disrupted by viruses. Here, we report Bcl-2 associated transcription factor 1 (Bclaf1) is degraded during the alphaherpesvirus Pseudorabies virus (PRV) and Herpes simplex virus type 1 (HSV-1) infections through the viral protein US3. We further reveal that Bclaf1 functions critically in type I interferon signaling. Knockdown or knockout of Bclaf1 in cells significantly impairs interferon-α (IFNα)-mediated gene transcription and viral inhibition against US3 deficient PRV and HSV-1. Mechanistically, Bclaf1 maintains a mechanism allowing STAT1 and STAT2 to be efficiently phosphorylated in response to IFNα, and more importantly, facilitates IFN-stimulated gene factor 3 (ISGF3) binding with IFN-stimulated response elements (ISRE) for efficient gene transcription by directly interacting with ISRE and STAT2. Our studies establish the importance of Bclaf1 in IFNα-induced antiviral immunity and in the control of viral infections.

## Introduction

Herpesviridae is a family of large DNA viruses with an ability to establish persistent infection in hosts. The viruses have evolved multiple strategies to establish persistent infection and combat host defenses; among these, the interferon (IFN) antiviral response is most prominent. Members of the family are causative agents of a variety of human and animal diseases and are further grouped into the three subfamilies, including alpha-, beta- and gammaherpesviruses (Steiner & Benninger, 2013). The alphaherpesvirus subfamily is neurotropic, including the genera simplexvirus and varicellovirus.

Pseudorabies virus (PRV) and herpes simplex virus type 1 (HSV-1) belong to the alphaherpesvirus subfamily and the genera varicellovirus and simplexvirus, respectively. They are often used as model viruses to study alphaherpesvirus biology. PRV is a swine pathogen that causes the economically important Aujeszky’s disease (Muller, Hahn et al., 2011, Pomeranz, Reynolds et al., 2005). HSV-1 is a human restricted virus, resulting in various mucocutaneous diseases, such as herpes labialis, genital herpes, herpetic whitlow, and keratitis (Roizman & Whitley, 2013). It also causes serious encephalitis in a small portion of the infected individuals (Roizman & Whitley, 2013).

Viral infection is defended by hosts at multiple levels, including intrinsic, innate and adaptive immunity. The type I Interferon (IFN-I) response plays a central role in innate immunity against viral infection. IFN-I positions cells in a potent antiviral state by inducing the synthesis of hundreds of antiviral proteins encoded by IFN-stimulated genes (ISGs). This process is initiated by binding of IFN-I to its receptor subunits (IFNAR1 and IFNAR2), which leads to the activation of the Janus Kinases (JAKs), JAK1 and TYK2. Activated JAKs then phosphorylate signal transducer and activator of transcription (STAT) 1 and 2, leading to the formation of a trimeric complex, referred to as IFN-stimulated gene factor 3 (ISGF3), which is comprised of STAT1/STAT2 and IFN regulatory factor 9 (IRF9). ISGF3 translocates to the nucleus and binds to IFN-stimulated response elements (ISRE) in the DNA to initiate the transcription of ISGs (Platanias, 2005, Stark & Darnell, 2012, Wang, Xu et al., 2017). Many of the gene products have potent antiviral functions (Sadler & Williams, 2008). Viruses have, in turn, evolved various strategies to antagonize the functions of IFN, which might be particularly important for herpesviruses to establish persistent infection in hosts (Garcia-Sastre, 2017, Katze, He et al., 2002, Schulz & Mossman, 2016). Key molecules in IFN signaling are targeted by various components of alphaherpesviruses. For example, PRV or HSV-1 utilize their encoded dUTPase UL50 to induce IFNAR1 degradation and inhibit type I IFN signaling in an enzymatic activity-independent manner (Zhang, Xu et al., 2017).

Increasing evidence indicates that IFN signaling is subject to extensive regulation and that additional coregulators are required to modulate the transcription of ISGs. For instance, the methyltransferase SETD2 promotes IFNα-dependent antiviral immunity via catalyzing STAT1 methylation on K525 (Chen, Liu et al., 2017); RNF2 increases the K33-linked polyubiquitination of STAT1 at position K379 to promote the disassociation of STAT1/STAT2 from DNA and suppress the transcription of ISGs (Liu, Jiang et al., 2018). The molecules that participate in IFN-induced transcription could be potential targets of herpesviruses. Thus, identifying novel components in IFN signaling and their interactions with viral molecules will provide a deeper understanding of IFN signaling and its interaction with viral infection.

US3 is a conserved Ser/Thr kinase encoded by every alphaherpesvirus identified thus far (Deruelle & Favoreel, 2011). It critically participates in the pathogenicity of viruses *in vivo* and is involved in the nuclear egress of viral capsids (Reynolds, Wills et al., 2002, Wagenaar, Pol et al., 1995). As a viral kinase, US3 expression impacts host cells in many aspects, including cytoskeletal alteration (Broeke, Radu et al., 2009, Favoreel, Minnebruggen et al., 2005, Jacob, Van den Broeke et al., 2015), the inhibition of histone deacetylase 1 and 2 (HDAC1/2) (Poon, Gu et al., 2006, Walters, Kinchington et al., 2010), and, more notably, disruption of various host defense mechanisms. US3 prevents host cells from apoptosis (Benetti & Roizman, 2007, Chang, Lin et al., 2013, Leopardi, Sant et al., 1997), disrupts the antiviral subnuclear structures promyelocytic leukemia nuclear bodies (PML-NBs) (Jung, Finnen et al., 2011), down-regulates major histocompatibility complex (MHC) class I surface expression (Rao, Pham et al., 2011), and interferes with the IFN response (Liang & Roizman, 2008, Piroozmand, Koyama et al., 2004, Wang, Ni et al., 2014, Wang, Wang et al., 2013).

Bclaf1 (Bcl-2 associated transcription factor 1; also called Btf for Bcl-2 associated transcription factor) was initially identified in a yeast two-hybrid system as a binding protein for adenovirus E1B 19K protein (Kasof, Goyal et al., 1999). It contains homology to the basic zipper (bZip) and Myb domains and binds DNA *in vitro* (Kasof et al., 1999). Bclaf1-knockout mice are embryonic lethal due to defects in lung development (McPherson, Sarras et al., 2009). Bclaf1 participates in diverse biological processes, including apoptosis (Kasof et al., 1999), autophagy (Lamy, Ngo et al., 2013), DNA damage response (Lee, Yu et al., 2012, Savage, Gorski et al., 2014), senescence (Shao, Sun et al., 2016), cancer progression (Dell’Aversana, Giorgio et al., 2017, Zhou, Li et al., 2014) and T cell activation (Kong, Kim et al., 2011). Recently, a role for Bclaf1 in herpesviral defense is emerging, and more strikingly, Bclaf1 is targeted by multiple viral components. The betaherpesviruse human cytomegalovirus (HCMV) dispatches both viral proteins (pp71 and UL35) and a microRNA to diminish cellular Bclaf1 levels (Lee, Kalejta et al., 2012). Bclaf1 is also identified as a target of several latently expressed microRNAs of the gammaherpesviruse Kaposi’s sarcoma-associated herpesvirus (KSHV) (Ziegelbauer, Sullivan et al., 2009). The fact that multiple mechanisms have been utilized by the members of beta- and gammaherpesviruses to suppress the expression of Bclaf1 indicates that this protein has a very important antiviral function. However, whether Bclaf1 is also involved in alphaherpesvirus infection and the molecular mechanism of its antiviral function are not known.

In this study, we examined the role of Bclaf1 in alphaherpesvirus infection and found that Bclaf1 is also degraded during PRV and HSV-1 infection through US3. More importantly, we revealed Bclaf1 as a critical regulator in the IFN-induced antiviral response. On the one hand, Bclaf1 maintains a mechanism that allows STAT1/STAT2 to be efficiently phosphorylated in response to IFN; on the other hand, it interacts with ISGF3 complex in the nucleus mainly through STAT2 and facilitates their interactions with the promoters of ISGs. These results reveal a critical role for Bclaf1 in IFN signaling and a strategy employed by alphaherpesvirus to disable it.

## Results

### PRV and HSV-1 dispatch US3 to degrade Bclaf1 in a proteasome-dependent manner

To examine the effect of alphaherpesvirus infection on Bclaf1, we infected porcine cells with PRV and human cells with HSV-1. We observed a dramatic decrease in Bclaf1 levels in all the cells examined at the time points when substantial viral proteins were expressed, including porcine kidney PK15 (Figure 1A), swine testis (ST) (Figure S1A) cells and human HEp-2 (Figure 1B) cells. Bclaf1 reduction appeared to occur more rapidly during HSV-1 infection. Since Bclaf1 is degraded in the proteasome upon HMCV infection, we examined if this was the case for PRV and HSV-1. We treated the cells with the proteasome inhibitor MG132 for 8 h at 1 h after viral adsorption. Compared with the control, the MG132 treatment blocked PRV and HSV-1 infection induced down-regulation of Bclaf1 and had minimal effect on viral protein expression (Figure 1C and 1D). These results suggest that both PRV and HSV-1 infection trigger a targeted and proteasome-dependent degradation of Bclaf1.

**Figure 1.**
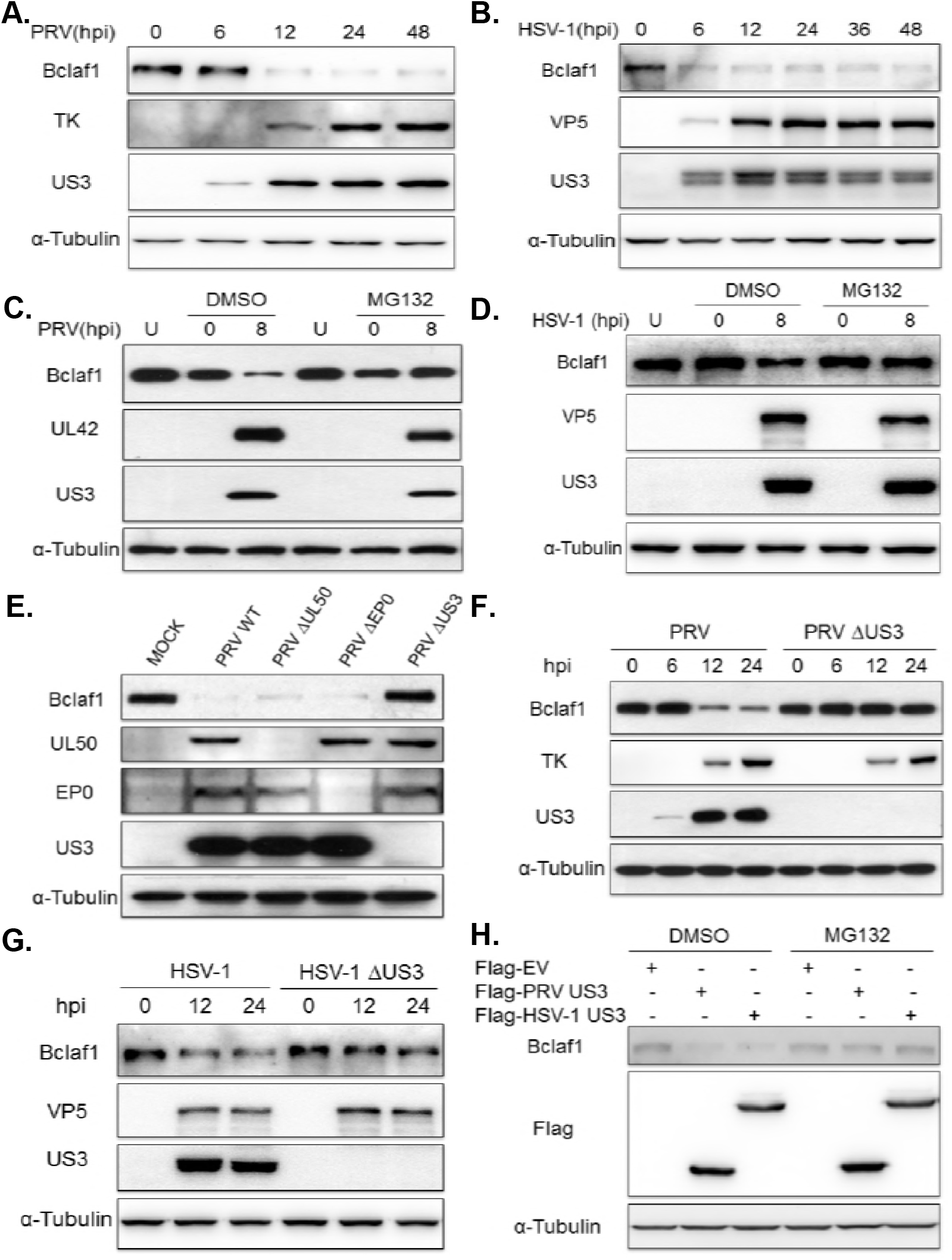
PRV and HSV-1 employ US3 to decrease Bclaf1 in a proteasome-dependent manner. (A) IB analysis of Bclaf1, TK and US3 in PK15 cells infected with PRV (MOI=1) for the indicated hours. α-Tubulin was used as loading control. (B) IB analysis of Bclaf1, VP5 and US3 in HEp-2 cells infected with HSV-1 (MOI=5) for the indicated hours. (C) IB analysis of Bclaf1, UL42 and US3 in PK15 cells infected with PRV (MOI=1) followed by untreatment (U) or treatment with DMSO or MG132. (D) IB analysis of Bclaf1, VP5 and US3 in HEp-2 cells infected with HSV-1 (MOI=5) followed by untreatment (U) or treatment with DMSO or MG132. (E) IB analysis of Bclaf1, UL50, EP0 and US3 in PK15 cells infected with indicated PRV strains (MOI=1) at 8 hpi. (F) IB analysis of Bclaf1, TK and US3 in PK15 cells infected with PRV WT or PRV ΔUS3 (MOI=1) for the indicated hours. (G) IB analysis of Bclaf1, VP5 and US3 in HEp-2 cells infected with HSV-1 WT or HSV-1 ΔUS3 (MOI=5) for the indicated hours. (H) IB analysis of endogenous Bclaf1 in HEK293T cells transfected with Flag-tagged PRV/HSV-1 US3 expression plasmids followed by treatment with DMSO or MG132.

To determine the viral protein responsible for the Bclaf1 degradation, we utilized a panel of gene deletion PRVs, particularly EP0, US3 and UL50 deleted strains, since these viral proteins are involved in the degradation of various proteins (Boutell & Everett, 2013, Jung et al., 2011, Zhang et al., 2017). Infecting cells with WT and the gene deletion PRVs showed that only the PRV ΔUS3 strain lost the ability to degrade Bclaf1 (Figure 1E). Indeed, although the Bclaf1 levels in the cells infected with PRV WT decreased over time up to 24 h post infection, those in the PRV ΔUS3 infected cells remained unchanged in the PK15 cells (Figure 1F) and even increased in the ST cells (Figure S1B, S1C and S1D). Similarly, the deletion of US3 from HSV-1 also abolished its ability to decrease Bclaf1 in the HEp-2 cells (Figure 1G). Collectively, these data indicate that US3 is essential for PRV- and HSV-1-induced Bclaf1 down-regulation. It also suggests that certain cells may respond to PRV and HSV-1 infection by increasing Bclaf1, which is concealed by US3 mediated Bclaf1 down-regulation.

To determine if US3 alone is sufficient to induce Bclaf1 degradation, we ectopically expressed PRV or HSV-1 US3 in HEK293T cells. The expression of US3 but not the empty vector or UL50 markedly reduced endogenous Bclaf1 (Figure S1E), which was rescued by MG132 treatment (Figure 1H). These results suggest that US3 induces the proteasomal degradation of Bclaf1.

### Bclaf1 promotes the IFNα-mediated inhibition of PRV/HSV-1 replication

The degradation of Bclaf1 upon PRV/HSV-1 infection by US3 suggests that Bclaf1 may possess an important antiviral function, which is inhibited by US3 but should be in action against US3 deficient viruses. Thus, to determine the role of Bclaf1 in viral infection, we focused on the differential properties between WT and ΔUS3 PRV infected cells. Although one well-known function of US3 is antiapoptosis, and Bclaf1 has been shown to be involved in it, we observed a similar level of apoptosis induced by ΔUS3 PRV infection in the Bclaf1 knockdown and control cells (Figure S2).

The dramatic difference we observed between the WT and ΔUS3 PRV/HSV1 was that the latter was more susceptible to interferon. The deletion of US3 in PRV/HSV-1 significantly decreased viral productions in PK15 (PRV) and HEp-2 (HSV-1) cells treated with IFNα, while having no or a slight influence on viral growth in the absence of interferon treatment (Figure 2A and 2B). As expected, Bclaf1 was not degraded in ΔUS3 PRV/HSV-1 infected cells. To determine whether Bclaf1 was involved in interferon mediated viral suppression, we depleted Bclaf1 using siRNAs in PK15 and HEp-2 cells or utilized a Bclaf1 knockout HeLa cell line and then infected the cells with ΔUS3 PRV/HSV-1 treated with or without IFNα. Compared with their respective controls, the expression of viral proteins and viral productions in Bclaf1 knockdown or knockout cells was significantly increased when treated with IFNα (Figure 2C, 2D and 2E). Altogether, our data supports that Bclaf1 enhances the IFNα-induced antiviral function against ΔUS3 PRV/HSV-1 infection.

**Figure 2.**
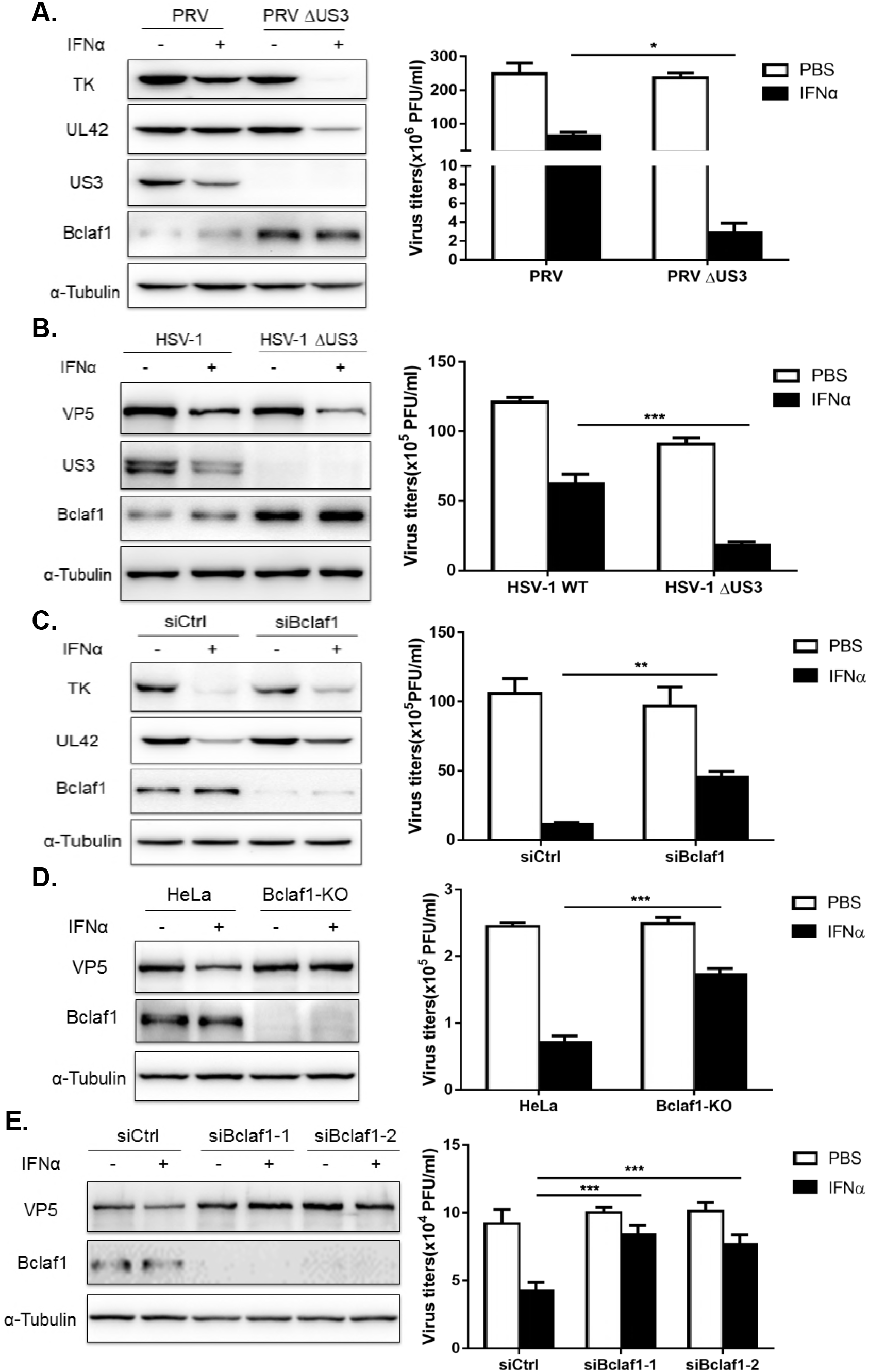
Bclaf1 contributes to the inhibition of IFNα to PRV and HSV-1. (A) PK15 cells were treated with PBS or porcine IFNα (500U/ml) for 12 hours followed infected with PRV WT or PRV ΔUS3 (MOI=0.5) for 24 hours. IB analyzed TK, UL42, US3 and Bclaf1 expression and plaque assay analyzed virus titers in supernatants. (B) HEp-2 cells were treated with PBS or human IFNα (500U/mL) for 12 hours followed infected with HSV-1 WT or HSV-1 ΔUS3 (MOI=1) for 24 hours. IB analyzed VP5, US3 and Bclaf1 expression and plaque assay analyzed virus titers in supernatants. (C) IB analysis of TK, UL42 and Bclaf1 in PK15 cells transfected with si-control or si-Bclaf1 followed by PBS or porcine IFNα (500U/mL) treatment for 12h and then infected with PRV ΔUS3 (MOI=1) for 24h. Plaque assay analyzed titers of virus in supernatants. (D) IB analysis of VP5 and Bclaf1 in control and Bclaf1-KO HeLa cells pre-treated with PBS or human IFNα (500U/mL) for 12h followed by HSV-1 ΔUS3 infection (MOI=5) for 24h. Plaque assay analyzed titers of virus in supernatants. (E) IB analysis of VP5 and Bclaf1 in HEp-2 cells transfected with si-control or si-Bclaf1 followed by PBS or human IFNα (500U/mL) treatment for 12h and then infected with HSV-1 ΔUS3 (MOI=3) for 24h. Plaque assay analyzed titers of virus in supernatants. Data are shown as mean ± SD of three independent experiments. Statistical analysis was performed by the two-way ANOVA test. *p<0.05; **p<0.01; ***p<0.001

### Bclaf1 is required for IFNα-induced ISG expression

We then examined the effect of Bclaf1 depletion on IFNα-induced gene transcription. Using an ISRE luciferase reporter assay, real time PCR and Western analysis, we showed that the IFNα-induced luciferase activity and upregulation of mRNAs and proteins of the examined ISGs were all much lower in HeLa Bclaf1-KO cells than those in control HeLa cells (Figure 3A, 3B and 3C). Knockdown of Bclaf1 in HEp-2 cells (Figure 3D) or in PK15 cells (Figure S3) using siRNAs also reduced IFNα-induced transcription. The deficiency of ISG induction in Bclaf1-KO HeLa cells after IFNα treatment was partially restored by the overexpression Bclaf1 (Figure 3E). Collectively, these data suggest that Bclaf1 enhances IFNα-induced transcription.

**Figure 3.**
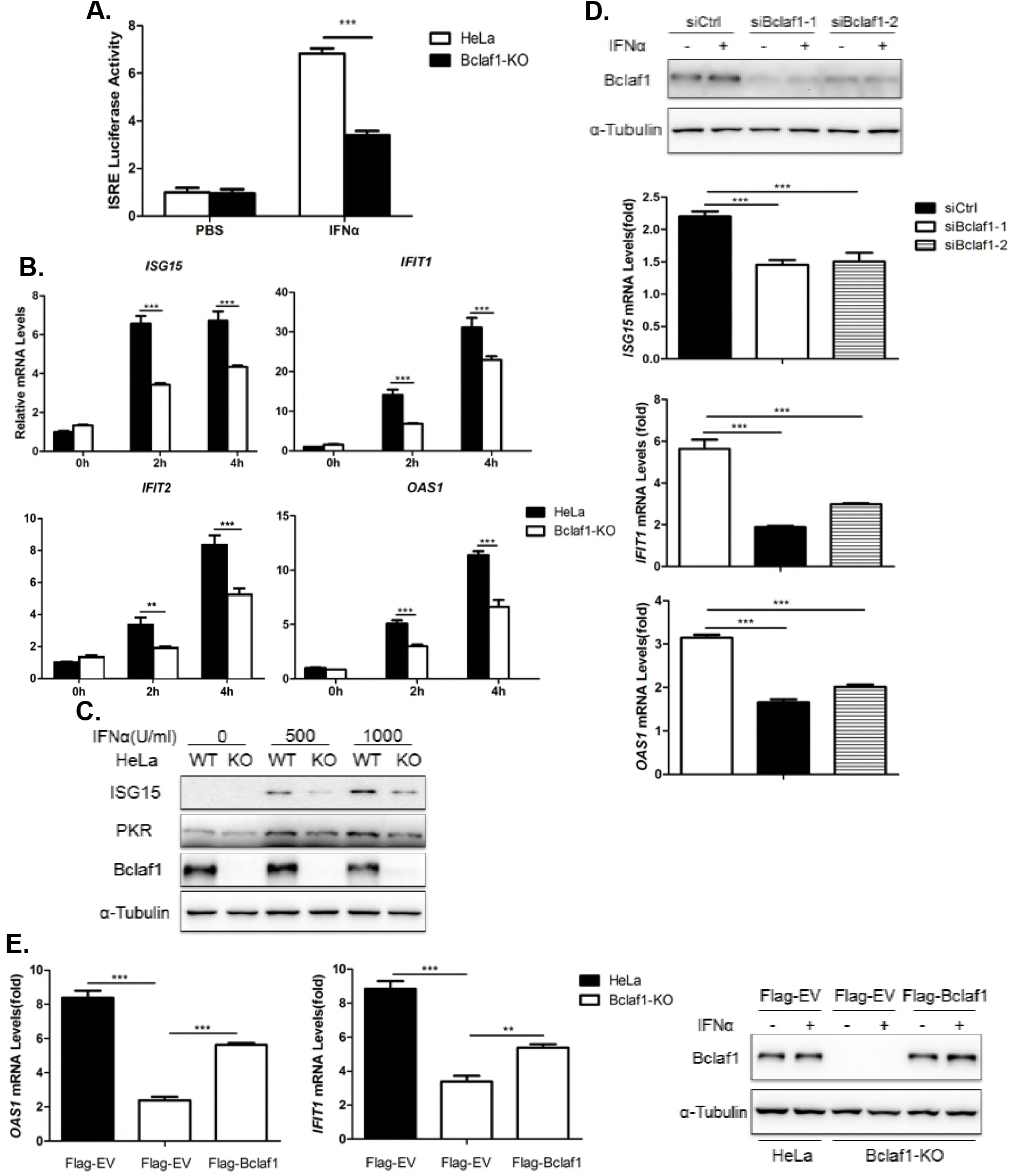
Bclaf1 facilitates IFNα-induced ISG expression. (A) ISRE-luciferase assay in HeLa WT and HeLa Bclaf1-KO cells treated with human IFNα (500U/mL) for 10h. (B) qRT-PCR analysis of *ISG15, IFIT1, IFIT2* and *OAS1* mRNA levels in HeLa WT and HeLa Bclaf1-KO cells treated with human IFNα (500U/mL) for the indicated time. (C) IB analysis of ISG15 and PKR in HeLa WT and HeLa Bclaf1-KO cells treated with human IFNα for 12h. (D) qRT-PCR analysis of *ISG15, IFIT1* and *OAS1* mRNA levels in HEp-2 cells transfected with si-control or si-Bclaf1 followed by human IFNα (500U/mL) treatment for 4h. IB analyzed the knocking down efficiency. (E) qRT-PCR analysis of *OAS1* and *IFIT1* mRNA levels in indicated HeLa cells transfected Flag-tagged EV or Bclaf1 expression plasmids followed by PBS or human IFNα (500U/mL) treatment for 4h. IB analyzed the expression of Bclaf1. Data are shown as mean ± SD of three independent experiments. Statistical analysis was performed by the two-way ANOVA test (A and B) and one-way ANOVA test (D and E). **p<0.01; ***p<0.001

### Bclaf1 facilitates the phosphorylation of STAT1/STAT2

To understand the exact role of Bclaf1 in the IFN signaling, we analyzed the signaling events that might be impaired in Bclaf1-deficient cells. We observed reduced courses of phosphorylation for STAT1 and STAT2 in response to IFNα in Bclaf1-KO HeLa cells (Figure 4A) and Bclaf1-silenced HEp-2 cells (Figure 4B) compared with relative control cells. Fractionation experiments demonstrated that the IFNα-induced nuclear translocation of STAT1/STAT2 in the Bclaf1-knockdown cells was reduced accordingly (Figure 4C). Thus, the loss of Bclaf1 impairs the IFNα-induced phosphorylation of STAT1/STAT2.

**Figure 4.**
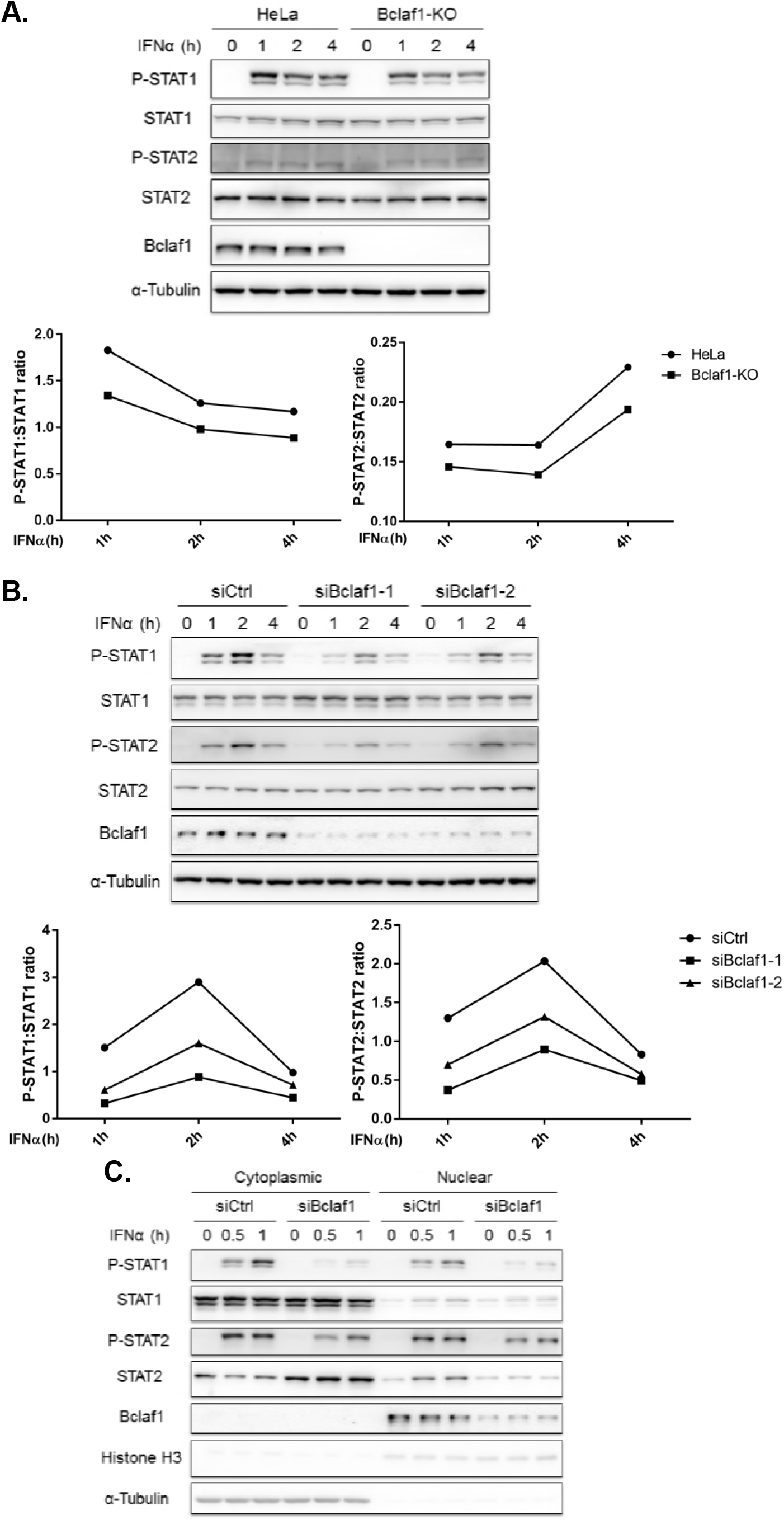
Loss of Bclaf1 attenuates IFNα-mediated STAT1/STAT2 phosphorylation. (A) IB analysis of phosphorylated(P)-STAT1, P-STAT2, STAT1, STAT2 and Bclaf1 in HeLa WT and HeLa Bclaf1-KO cells treated with human IFNα (500U/mL) for the indicated time. Data were quantified and shown as the ratio of P-STAT1 to STAT1 and P-STAT2 to STAT2. (B) IB analysis of P-STAT1, P-STAT2, STAT1, STAT2 and Bclaf1 in HEp-2 cells transfected with si-control or si-Bclaf1 followed by PBS or human IFNα (500U/mL) treatment for the indicated time. Data were quantified and shown as the ratio of P-STAT1 to STAT1 and P-STAT2 to STAT2. (C) IB analysis of P-STAT1, P-STAT2, STAT1, STAT2 and Bclaf1 in cytoplasmic and nuclear extracts of HEp-2 cells transfected with si-control or si-Bclaf1 followed by PBS or human IFNα (500U/mL) treatment for the indicated time. α-Tubulin and Histone H3 were used as the cytoplasmic and nuclear controls, respectively.

Because the majority of Bclaf1 localized in the nucleus, the mechanism for Bclaf1 to influence this step is likely indirect, possibly through altering the expression levels of the components essential for STAT1/STAT2 phosphorylation. However, no obvious difference in the major components, including Receptor, JAK1, TYK2, STAT1 and STAT2, between the Bclaf1 knockdown or knockout cells and the WT controls was observed (Figure S4).

### Bclaf1 binds with ISRE and promotes the association of ISGF3 with DNA

In addition, our Chromatin Immunoprecipitation (ChIP) assays showed that IFNα-induced binding of STAT1/STAT2 to the promoters of ISGs was also greatly decreased in Bclaf1-KO HeLa cells (Figure 5A) and Bclaf1-silenced HEp-2 cells (Figure S5) compared with that in relative control cells.

**Figure 5.**
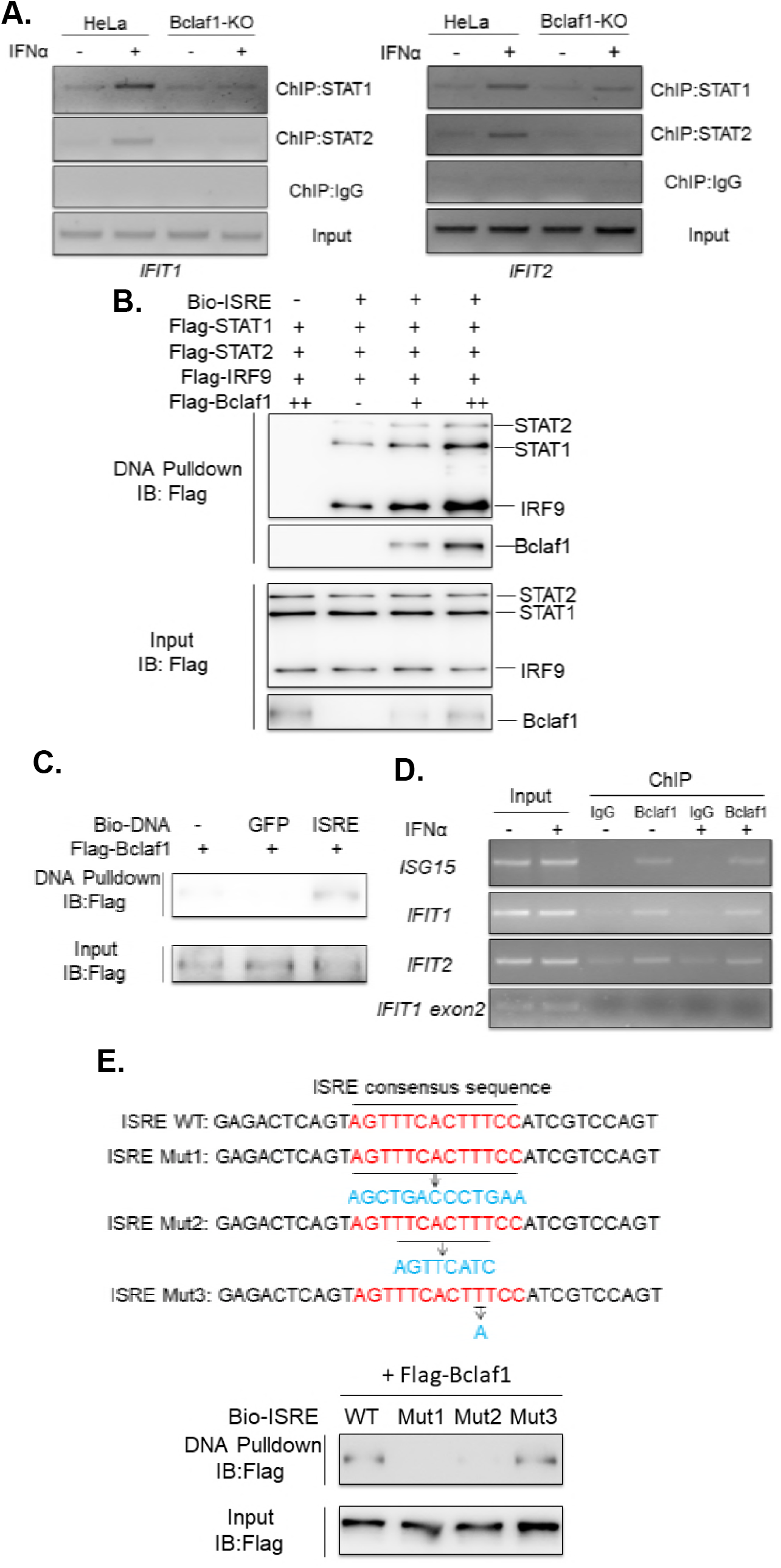
Bclaf1 binds with ISRE and promotes the association of ISGF3 with DNA. (A) ChIP analysis of STAT1/STAT2 DNA-binding in promoters of *IFIT1* and *IFIT2* in HeLa WT and HeLa Bclaf1-KO cells simulated with PBS or human IFNα (500U/mL) for 1h. (B) IB analysis of Bio-ISRE pull-down STAT1, STAT2, IRF9 and Bclaf1. Unlabeled ISRE was used for control. (C) IB analysis of ISRE-binding Bclaf1. Unlabeled ISRE and Bio-GFP were used for control. (D) ChIP analysis of Bclaf1 DNA-binding in promoters of *ISG15, IFIT1* and *IFIT2* in HeLa cells simulated with PBS or human IFNα (500U/mL) for 1h. An amplicon located in *IFIT1 exon2* was also tested for control. (E) IB analysis of WT or mutated (1–3) Bio-ISRE pull-down Bclaf1.

Because Bclaf1 predominantly localized in the nucleus, we reasoned that Bclaf1 should exert its function in the nucleus and that the reduced STAT1/STAT2 phosphorylation by IFNα upon Bclaf1 reduction could be an indirect consequence. Therefore, we focused on the aspect that Bclaf1 may enhance the binding of ISGF3 to the promoters. To exclude the possibility that the impaired binding between STAT1/STAT2 to the ISGs promoters in the Bclaf1-knockdown cells was due to the reduced nuclear STAT1/STAT2 in these cells, we performed a DNA pull-down assay to directly measure whether STAT1/STAT2/IRF9 binding to the promoters was enhanced by Bclaf1. An ISRE DNA was synthesized, biotin-labeled, and added into equal amounts of purified STAT1/STAT2/IRF9 as well as increased concentrations of purified Bclaf1 followed by a streptavidin-bead pull-down. The addition of Bclaf1 drastically increased the binding of STAT1/STAT2/IRF9 to Bio-ISRE in a dose-dependent manner, and Bclaf1 was present in the Bio-ISRE pull-down complex (Figure 5B). Purified Bclaf1 was pulled down by Bio-ISRE but not by Bio-GFP (Figure 5C), suggesting that Bclaf1 was directly bound to ISRE specifically. The ChIP assay confirmed that Bclaf1 was bound to the promoter regions of ISGs in HeLa cells (Figure 5D), which appeared to be constitutive and was not induced by IFNα treatment. To further characterize the DNA sequence required for binding with Bclaf1, we replaced entire ISRE consensus sequence (Mut1) or the core sequence of 5’-TTCNNTTT-3’ (Au-Yeung, Mandhana et al., 2013) (Mut2) with a sequence from GFP. We also mutated the TTT motif near the 3’ end of the ISRE by chancing TTT to TAT (Mut3). DNA pull-down assays demonstrated that Mut1 and Mut2 failed to interact with Bclaf1, whereas Mut3 still could (Figure 5E), indicating Bclaf1 binds with the core sequence of ISRE and the TTT motif is not required. In aggregate, these data demonstrated that Bclaf1 bound with ISRE specifically and promoted the association of ISGF3 with DNA.

### Bclaf1 associates with ISGF3

To understand the molecular mechanism by which Bclaf1 facilitates ISGF3 binding to ISGs promoters, we performed co-IP assays to examine the interaction between Bclaf1 and ISGF3, which is composed of STAT1, STAT2 and IRF9. We constructed a HEp-2 cell line that endogenously expresses Flag-Bclaf1 by adding a *Flag* to the *Bclaf1* gene using the CRISPR/Cas9 technique and is referred as HEp-2-Flag-Bclaf1. Fractionation of the cells followed by co-IPs using a Flag-antibody showed that Flag-Bclaf1 interacted with STAT1, STAT2 and IRF9 in the nucleus where it mainly localized (Figure 6A and 6B). Reversely, endogenous Bclaf1 was also detected in the immuno-complexes of STAT1, STAT2 or IRF9 after IPs of nuclear extracts of HeLa cells using their respective antibodies (Figure S6). The interaction between Bclaf1 and STAT1/STAT2/IRF9 occurred in the absence of IFNα treatment and was increased after IFNα treatment, correlating with more STAT1/STAT2/IRF9 being translocated into the nucleus (Figure 6A, 6B and Figure S6). We further determined the regions in Bclaf1 that mediated its association with STAT1, STAT2 or IRF9 by co-expressing various Flag tagged Bclaf1 fragments with Ha tagged STAT1, STAT2 or IRF9 in HEK293T cells and performing co-IPs, and identified the region 236–620 responsible for binding to these proteins (Figure 6C). To examine whether the interaction between Bclaf1 and STAT1/STAT2/IRF9 is required for its ability to enhance IFNα transcription, we overexpressed Bclaf1 full-length and the indicated fragments in HEp-2 followed by IFNα treatment. mRNA measurements showed that the IFNα-induced *IFIT1* transcription was enhanced by full-length Bclaf1 and Bclaf1-F2 (236–620), and not by the fragments that failed to bind with STAT1/STAT2/IRF9 (Figure 6D). Taken together, these results suggest that Bclaf1 interacts with ISGF3 complex in the nucleus, which is important for Bclaf1 to enhance the activation of ISRE after IFNα stimulation.

**Figure 6.**
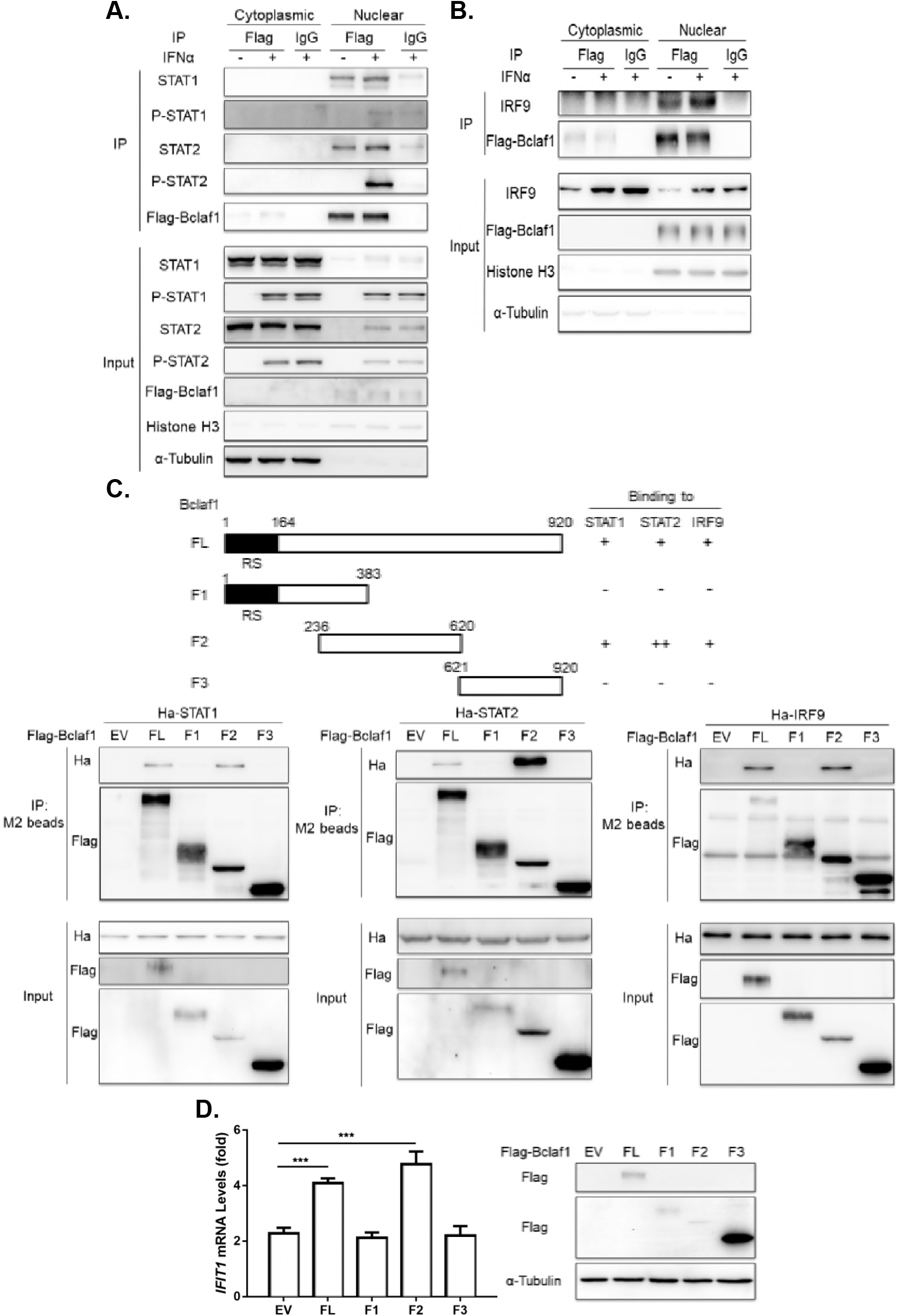
Bclaf1 interacts with STAT1/STAT2/IRF9. (A) IB analysis of STAT1, STAT2, P-STAT1, P-STAT2 and Flag-Bclaf1 in cytoplasmic or nuclear immunoprecipitates of a HEp-2-Flag-Bclaf1 cell line treated with PBS or human IFNα (500U/mL) for 2h. IgG was used for control immunoprecipitation. (B) IB analysis of IRF9 and Flag-Bclaf1 in cytoplasmic or nuclear immunoprecipitates of a HEp-2-Flag-Bclaf1 cell line treated with PBS or human IFNα (500U/mL) for 4h. (C) IB analysis of immunoprecipitates of HEK293T cells co-transfected with Flag-tagged Bclaf1 truncations and Ha-tagged STAT1/STAT2IRF9 expression plasmids. (D) qRT-PCR analysis of *IFIT1* mRNA levels in HEp-2 cells transfected with Flag-tagged EV, full-length Bclaf1 or its truncations expression plasmids followed by PBS or human IFNα (500U/mL) treatment for 3h. IB analyzed the expression of Bclaf1. Data are shown as mean ± SD of three independent experiments. Statistical analysis was performed by the one-way ANOVA test. ***p<0.001

### Bclaf1 associates with ISGF3 complex primarily through interacting with STAT2

Next, we set out to determine how Bclaf1 interacts with ISGF3. We first examined the direct interactions between Bclaf1 and the components of ISGF3 by mixing bacterially purified His-STAT1, -STAT2 or -IRF9 with GST-Bclaf1 F2 followed by GST pull-down assays. Western analysis showed that only His-STAT2 was able to be pulled down specifically by GST-Bclaf1 F2, whereas the other two were not (Figure 7A and data not shown). These results hinted that STAT2 is the crucial component connecting ISGF3 to Bclaf1. In supporting this, co-IP assays showed that the interaction between Bclaf1 and STAT1 or IRF9 was enhanced by STAT2, and not by IRF9 or STAT1 upon overexpression in 293T cells (Figure 7B and 7C). Moreover, the interaction between Bclaf1 and STAT1 or IRF9 at endogenous levels was decreased upon STAT2 knockdown in HEp-2-Flag-Bclaf1 cells treated with IFNα (Figure 7D). In addition, *in vitro* DNA pulldown assays demonstrated that in the absence of STAT2 Bclaf1 lost its ability to recruit the components of ISGF3 to ISRE (Figure 7E). Collectively, these data indicate that STAT2 is the key component mediating the binding of Bclaf1 to ISGF3 complex.

**Figure 7.**
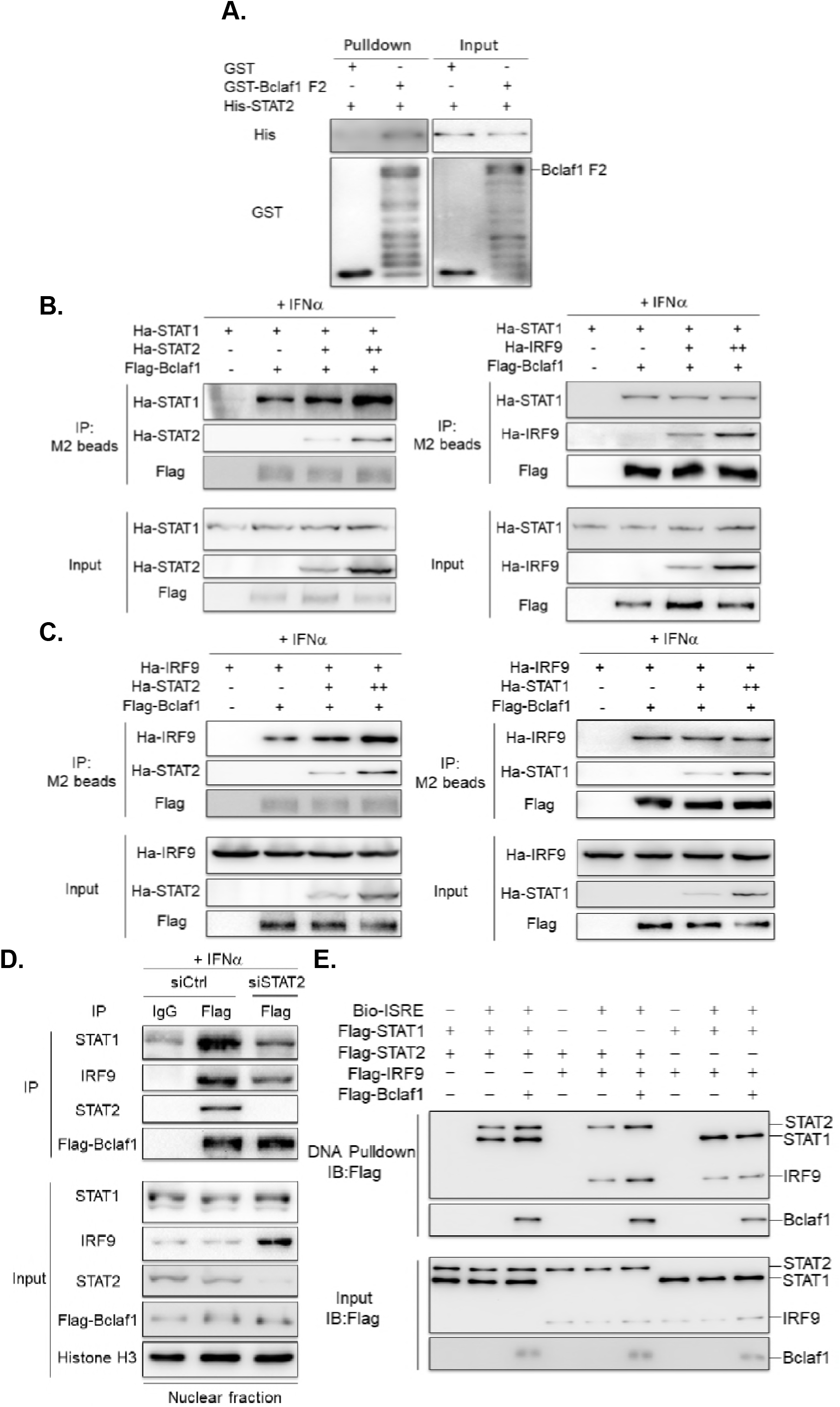
Bclaf1 interacts with ISGF3 mainly through STAT2. (A) GST pulldown analysis of the interaction between His-STAT2 and GST-Bclaf1 F2. (B) IB analysis of immunoprecipitates of HEK293T cells co-transfected with Flag-tagged Bclaf1, Ha-tagged STAT1 or STAT2/IRF9 expression plasmids. (C) IB analysis of immunoprecipitates of HEK293T cells co-transfected with Flag-tagged Bclaf1, Ha-tagged IRF9 or STAT2/STAT1 expression plasmids. (D) IB analysis of STAT1, STAT2, IRF9 and Flag-Bclaf1 in nuclear immunoprecipitates of a HEp-2-Flag-Bclaf1 cell line transfected with si-control or si-STAT2 followed by PBS or human IFNα (500U/mL) treatment for 3h. (E) IB analysis of Bio-ISRE pull-down STAT1, STAT2, IRF9 and Bclaf1.

## Discussion

The IFN response is critical in the control of viral infection and is often evaded or antagonized by various viruses. Most identified strategies used by viruses to evade ISG expression emphasize on the known signaling molecules in the IFN pathway targeted by various viral components. Here, we revealed a novel positive regulator, Bclaf1, in IFN signaling and its degradation by the viral protein US3 during alphaherpesvirus PRV and HSV-1 infection.

The evidence supporting Bclaf1 as a critical regulator in IFN-mediated antiviral response includes the following: 1) IFNα-induced ISG transcription is greatly compromised in Bclaf1 knockdown or knockout cells; 2) Bclaf1 is required for the efficient phosphorylation of STAT1 and STAT2 induced by IFNα; 3) Bclaf1 binds with ISRE and facilitates the binding of ISGF3 complex to promoters of the ISGs; 4) Bclaf1 interacts with ISGF3 through STAT2; 5) Bclaf1 is degraded by US3 during PRV and HSV-1 infection; and 6) In the absence of US3, PRV and HSV-1 become more sensitive to IFNα treatment, which is partly due to the unreduced level of Bclaf1 in the cells. These findings establish Bclaf1 as a critical positive regulator in IFN signaling and indicate its importance in host innate immunity against herpesvirus infection, which may be more broadly against other viruses as well.

We demonstrated that Bclaf1 was involved in two critical steps in IFN signaling, including the efficient phosphorylation of STAT1 and STAT2 and binding of the transcriptional complex to ISGs promoters (Figure 8). At present, the mechanism by which Bclaf1 regulates STAT1/STAT2 phosphorylation is unknown. STAT1/STAT2 phosphorylation is catalyzed by JAK1 and TYK2 activated by IFN-induced receptor dimerization, which occurs rapidly in the membrane. The mechanism for Bclaf1 to influence this step is likely indirect as Bclaf1 primarily localized in the nucleus. Emerging evidence indicates that the modification states of these components, prior to IFN engagement, also affect STAT1 and STAT2 phosphorylation by JAKs (Begitt, Droescher et al., 2011, Chen et al., 2017, Ginter, Bier et al., 2012, Liu et al., 2018, Steen, Nogusa et al., 2013, Wang, Nan et al., 2017). For instance, Chen et al showed that methyltransferase SETD2-mediated methylation of STAT1 significantly enhanced STAT1 phosphorylation by JAK1 (Chen et al., 2017). The result that the lack of Bclaf1 decreases STAT1/STAT2 phosphorylation without affecting the expression of upstream components suggests that Bclaf1 may be involved in pre-existing modifications of STAT1/STAT2 by regulating relevant enzymes.

**Figure 8.**
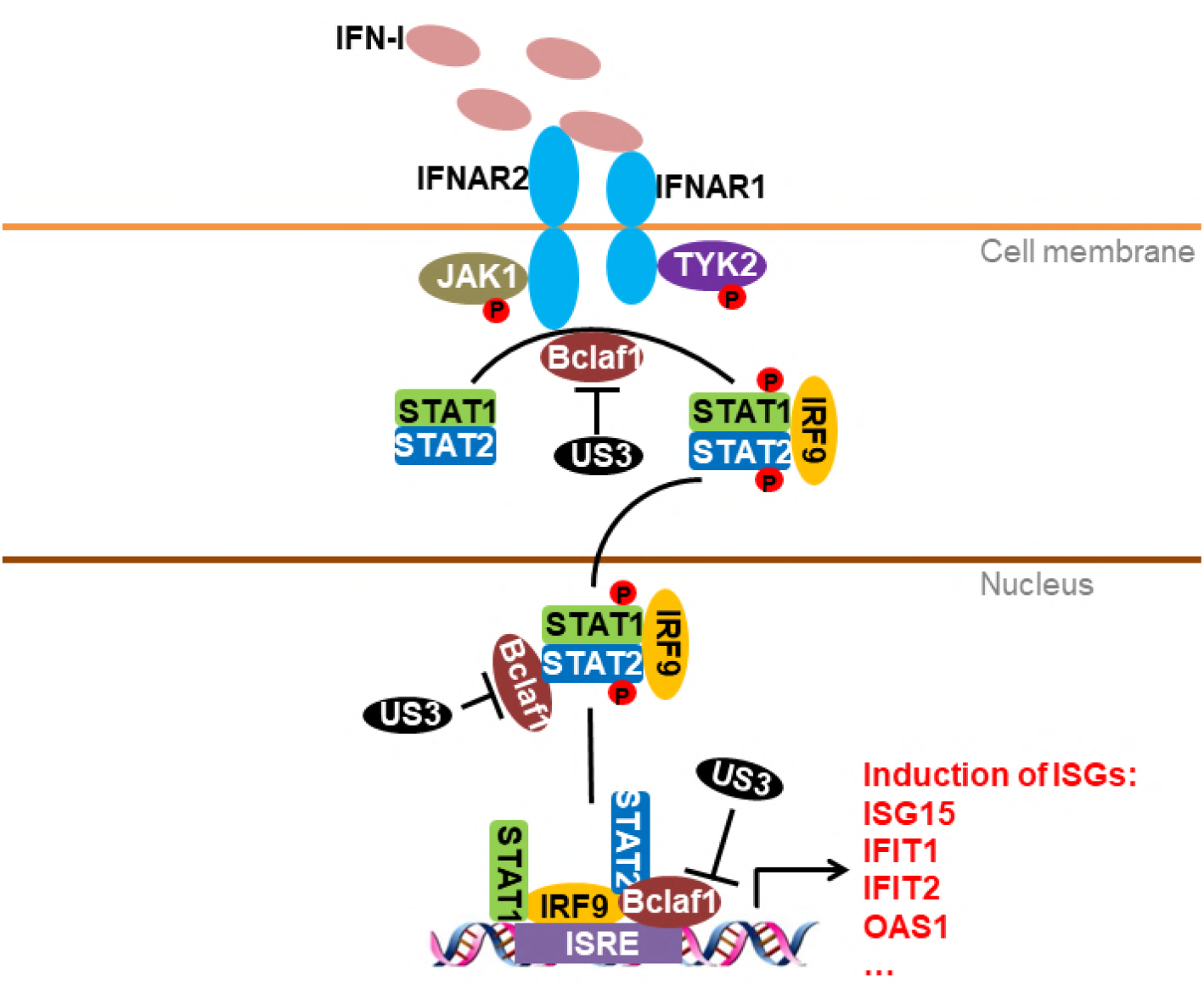
A working model of how Bclaf1 regulates IFN response pathway. Upon IFN-I binding with receptors, Bclaf1 facilitates the phosphorylation of STAT1/STAT2 in an indirect manner. Phosphorylated STAT1 and STAT2 associate with IRF9 to format a complex called ISGF3 and translocate to the nucleus. Bclaf1 is acting as a mediator attracting ISGF3 to ISGs promoters for efficient transcription. On the one hand, Bclaf1 interacts with STAT2 directly to associate with ISGF3. On the other hand, Bclaf1 binds with ISRE. During PRV or HSV-1 infection, US3 is dispatched to degrade Bclaf1 to inhibit IFN signaling.

Although the JAK-STAT pathway is well established, the regulation of the STAT1/STAT2/IRF9-mediated transcription of ISGs in the nucleus is not fully understood. We demonstrated that Bclaf1 is an important positive regulator in this process. Although epigenetic modifications and chromatin-remodeling, in the context of the promoter region, are important avenues for the regulation of transcription (Bonasio, Tu et al., 2010, Venkatesh & Workman, 2015), Bclaf1 appears to function by enhancing the recruitment of ISGF3 complex to the promoter of the ISGs by simultaneously binding to the promoter of the ISGs and this complex. Bclaf1 constitutively bound to the promoter of the ISGs without being enhanced by IFNα. It also interacted with ISGF3 in the nucleus, which was not regulated by IFNα-induced STAT1/STAT2 phosphorylation. However, as more and more STAT1/STAT2/IRF9 entered the nucleus following the IFNα treatment, more STAT1/STAT2/IRF9 was found to bind to Bclaf1 and the promoter of the ISGs as well. Thus, one conceivable role of Bclaf1 in ISGF3 mediated transcription is acting as a mediator attracting ISGF3 to its prebound ISGs promoters for efficient transcription. A similar mode of action is also observed in Bclaf1-regulated C/EBPβ transcription (Shao et al., 2016). Bclaf1 has a DNA-binding ability (Kasof et al., 1999), and we found that the binding between Bclaf1 and the promoter of the ISGs was likely to be a direct event. It would be interesting to further elucidate how Bclaf1 interacts with the promoter of the ISGs.

US3 is a potent alphaherpesviral kinase involved in antagonizing a wide range of host antiviral mechanisms. Here, we uncovered a strategy for US3 to impair IFN-mediated antiviral activity, which is to degrade Bclaf1. Bclaf1 was degraded by both genera of alphaherpeviruses and was also inhibited by members of beta- and gammaherpesviruses, indicating that the disruption of Bclaf1 might be a general mechanism for all herpesvirus infections. Since a key feature of herpesviruses is the establishment of a persistent infection and reactivation upon stress, Bclaf1 may participate in these processes. To establish persistent infection, herpesviruses employ multiple strategies to counteract the antiviral activity of IFN (Paladino & Mossman, 2009, Su, Zhan et al., 2016), and the disruption of Bclaf1 might be an integral part of sabotaging IFN signaling by herpesviruses. In addition, Bclaf1 possesses other antiviral functions, such as restriction of HMCV replication and inhibition of KSHV reactivation. Others and our studies have highlighted an important role of Bclaf1 against herpesviruses infection, and it may be broadly for other viruses as well. Thus, evaluating Bclaf1’s antiviral function *in vivo* is highly desirable. Because Bclaf1 knockout leads to embryonic lethality in mice, a conditional knockout mouse should be created.

## Materials and Methods

### Reagents

MG132 was purchased from APExBIO (133407-82-6). Streptavidin beads (3419) were purchased from Cell Signaling Technology. Flag M2 beads (A2220) and 3xFlag peptide (F4799) were purchased from Sigma. Human IFNα was purchased from PEPROTECH (300-02AA). Glutathione agarose was purchased from GE Healthcare (17-0756-01). Porcine IFNα was described previously (Zhang et al., 2017). Biotin 3’ End DNA Labeling Kit was purchased from Thermo Scientific (89818).

The following antibodies were used for co-Immunoprecipitation (co-IP): anti-Bclaf1 (1:100, sc-135845, Santa Cruz), anti-Flag (1:200, F1804, Sigma), anti-STAT1 (1:100, 14995, Cell Signaling Technology), anti-STAT2 (1:50, 72604, Cell Signaling Technology), and anti-IRF9 (1:50, 76684, Cell Signaling Technology). The following antibodies were used for Chromatin Immunoprecipitation (ChIP): anti-Bclaf1 (1:50, sc-135845, Santa Cruz), anti-STAT1 (1:50, 14995, Cell Signaling Technology) and anti-STAT2 (1:50, 72604, Cell Signaling Technology). The following antibodies were used for immunoblot analysis: anti-Bclaf1 (1:500, sc-135845, Santa Cruz), anti-Flag (1:2000, F1804, Sigma), anti-α-Tubulin (1:8000, PM054, MBL), anti-HA (1:1000, sc-805, Santa Cruz), anti-GFP (1:1000, sc-9996, Santa Cruz), anti-ISG15 (1:500, sc-166755, Santa Cruz), anti-PKR (1:1000, 12297, Cell Signaling Technology), anti-STAT1 (1:1000, 14995, Cell Signaling Technology), anti-STAT2 (1:1000, 72604, Cell Signaling Technology), anti-P-STAT1 (Tyr701) (1:1000, 9167, Cell Signaling Technology), anti-P-STAT2 (Tyr690) (1:1000, 88410, Cell Signaling Technology), anti-IRF9 (1:1000, 76684, Cell Signaling Technology), anti-JAK1 (1:500, 3344, Cell Signaling Technology), anti-TYK2 (1:1000, 14193, Cell Signaling Technology), anti-Histone H3 (1:2000, 17168-1-AP, Proteintech), and anti-caspase3 p17 (1:1000, sc-166589, Santa Cruz). The antibodies against PRV TK, PRV US3, PRV EP0, PRV UL50, and HSV-1 VP5 were described previously(Han, Chadha et al., 2012, Xu, Qin et al., 2015, Zhang et al., 2017). Mouse polyclonal antibodies against PRV UL42 and HSV-1 US3 were raised in mice individually with the N-terminal region of each protein as antigens.

### Cell and viruses

HEK293T cells (human embryonic kidney, ATCC #CRL-3216), HeLa cells (ATCC #CCL-2), HEp-2 cells (a kind gift from Dr. Xiaojia Wang which was described previously (Wang, Patenode et al., 2011)), PK15 cells (ATCC #CCL-33), ST cells (swine testis, ATCC #CRL-1746), and Vero cells (ATCC #CCL-81) were cultured in medium supplemented with 10% (v/v) FBS at 37°C and 5% CO2. The PRV Bartha-K61, recombinant PRV UL50-knockout virus (PRV ΔUL50), PRV EP0-knockout virus (PRV ΔEP0) and KOS strain of HSV-1 were described previously (Han et al., 2012, Xu et al., 2015). The recombinant PRV US3-knockout virus (PRV ΔUS3) and the HSV-1 US3-knockout virus (HSV-1 ΔUS3) were generated in this paper (see below).

### Plasmids

The PRV US3 gene was amplified from the Bartha-K61 genome, and the HSV-1 US3 gene was amplified from the KOS genome. Both PRV and HSV-1 US3 were cloned into the pRK5 vector with an N-terminal Flag tag. pRK5-Flag-PRV UL50, pRK5-Flag-HSV-1 UL50 and pRK5-Flag-Bclaf1 were previously described (Shao et al., 2016, Zhang et al., 2017). Bclaf1 truncations were amplified by PCR from pRK5-Flag-Bclaf1 and were cloned into the pRK5 vector with an N-terminal Flag vector. pRK5-Ha-STAT1/STAT2/IRF9 were constructed by amplifying STAT1/STAT2/IRF9 ORFs by PCR from cDNA synthesized from the total RNA of IFNα-stimulated HeLa cells and cloning it into the pRK5 vector with an N-terminal Ha tag vector.

### Real-Time PCR

Total RNA was extracted using TRIzol (Invitrogen) following the manufacturer’s protocol. A total of 0.8 μg total RNA from different treatments was reverse transcribed using M-MLV reverse transcriptase (Promega) with an oligo(dT) 18 primer. Real-time PCR was performed using an UltraSYBR Mixture (Beijing CoWin Biotech, Beijing, China) and a ViiA 7 real-time PCR system (Applied Biosystems). Sample data were normalized to GAPDH expression. Specific primers used for RT–PCR assays are listed in Supplementary Table 1.

### Immunoprecipitation and Western Blot

Cells were harvested and lysed in lysis buffer (50 mM Tris-Cl at pH 8.0, 150 mM NaCl, 1% Triton X-100, 1 mM DTT, 1× complete protease inhibitor cocktail tablet and 10% glycerol). The nuclear and cytoplasmic extracts from cells were prepared using a Nuclear and Cytoplasmic Protein Extraction Kit (Beyotime Biotechnology, Shanghai, China) following the manufacturer’s instructions. Equalized extracts were used for the immunoprecipitation and immunoblot analysis, which were described previously (Cui, Li et al., 2014).

### Luciferase Assay

Bclaf1-KO HeLa cells or control HeLa cells were seeded in 24-well plates and were then transfected with 100 ng of ISRE-luciferase reporter plasmids plus 20 ng of pRL-TK plasmids as an internal control. After 24 h of incubation, the cells were stimulated with PBS or IFNα, and whole-cell lysates were collected to measure the luciferase activity with a dual luciferase reporter assay kit (Promega).

### Virus Infection and Plaque Assay

PRV or HSV-1 were propagated and tittered in Vero cells. To infect, the cells were incubated with PRV or HSV-1 for 1 h, washed with PBS, and incubated in DMEM supplemented with 5% FBS until the times indicated. For the MG132 (ApexBio) treatment, a final concentration 20μM of MG132 was added into culture medium at 1 h post infection to allow efficient viral entry.

The Viral yield was determined by tittering in the Vero cells. Briefly, infected cell supernatants were cleared of cell debris by centrifugation. The Vero cells were infected in duplicate or triplicate with serial dilutions of supernatants for 1 h in serum free DMEM, washed with PBS, overlaid with 1× DMEM/1% agarose, and incubated at 37°C until plaque formation was observed (72 h-96 h). The cells were stained with 0.5% neutral red for 4 h-6 h at 37°C, and the plaques were counted.

### Generation of Recombinant PRV or HSV-1

PRV ΔUS3 was generated according to methods described previously (Xu et al., 2015). Briefly, PK15 cells were cotransfected with the viral genome and the CRISPR/Cas9 system containing two targeting sgRNAs for US3. After PRV-mediated CPE was prominently observed, the supernatants were collected, and the plaque assay was performed for subcloning the viruses. Single colonies were determined via sequencing and a Western blot with PRV US3 antibodies. For generation of HSV-1 ΔUS3, HEK293T cells were transfected using the CRISPR/Cas9 system containing the targeting sgRNA for US3, and 24 h later, the cells were infected with HSV-1 (KOS) at an MOI of 1. Viruses in the supernatants were collected at 48 h post infection and was subcloned via plaque assays. Single colonies were determined via sequencing and a Western blot with HSV-1 US3 antibodies.

Oligonucleotides used in this study are listed in Supplementary Table 1.

### Generation of Bclaf1-KO Cells

HeLa cells were seeded into a 6-well dish to achieve 70% confluency and were transfected with CRISPR/Cas9 plasmids containing a target sequence complimentary to the fourth exon of Bclaf1, and 48 h later, the cells were diluted and seeded into a 96-well dish at 0.5 cell/well in complete DMEM media. Wells that contained a single colony were expanded until enough cells were available for total protein extraction and determining Bclaf1 via a Western blot. Oligonucleotides used in this study are listed in Supplementary Table 1.

### Generation of a HEp-2 cell line that endogenously expresses Flag-Bclaf1

To add a Flag tag to the endogenous Bclaf1, HEp-2 cells were seeded into a 6-well dish to achieve 70% confluency and were transfected with CRISPR/Cas9 plasmids containing a target sequence complimentary to the intron that was prior to the ATG of Bclaf1 plus a donor plasmid containing homologous arms and Puro-P2A-3×Flag sequences. After 48 h, medium containing 2.5 mg/ml puromycin was added to select for tagged cells, and 48 h later, the cells were diluted and seeded into a 96-well dish at 0.5 cell/well in complete DMEM media. Wells that contained a single colony were expanded until enough cells were available for total protein extraction and determining Flag-Bclaf1 via a Western blot.

Oligonucleotides used in this study are listed in Supplementary Table 1.

### RNA Interference

siRNAs against Bclaf1 (1# 5’-GGTTCACTTCGTATCAGAA-3’) and (2# 5’-TTCTCAGAATAGTCCAATT-3’) and STAT2 (5’-CCCAGUUGGCUGAGAUGAUCUUUAA-3’) were transfected using Lipofectamine RNAiMax (Invitrogen) at a final concentration of 20 nM following the manufacturer’s instructions.

### Chromatin Immunoprecipitation (ChIP)

The ChIP assay was performed using a ChIP-IT Express enzymatic system (Active Motif, Carlsbad, CA, USA) following the manufacturer’s instructions. Briefly, cells were crosslinked with 1% formaldehyde and neutralized with 0.125 M glycine. Purified chromatin was digested to ~ 500 bp by enzymatic shearing. Anti-Bclaf1, anti-STAT1, anti-STAT2 or control IgG antibodies were used for immunoprecipitation. After reverse crosslinking, the DNA samples were analyzed by PCR followed by 3% agarose gel electrophoresis. Specific primers used are listed in Supplementary Table 1.

### DNA Pulldown assay

Flag-STAT1, Flag-STAT2, Flag-IRF9 and Flag-Bclaf1 were purified from overexpressed HEK293T cells stimulated with (STAT1/STAT2/IRF9) or without (Bclaf1) IFNα by immunoprecipitation using M2 beads (Sigma). The biotinylated ISRE (5’-GAGACTCAGTAGTTTCACTTTCCATCGTCCAGT-3’) DNA oligos were synthesized by a Biotin 3’ End DNA Labeling Kit (Thermo Scientific) and were then annealed and incubated with the purified indicated Flag-tagged proteins for 30 min in binding buffer (10 mM Tris, 1 mM KCl, 1%NP-40, 1 mM EDTA, 5% glycerol) at room temperature. Then, streptavidin beads (Cell Signaling) were added for incubation at 4°C for 1 h. After three washes with binding buffer, the ISRE-binding proteins were eluted by boiling and analyzed by immunoblotting.

### GST Pulldown

Purified His-STAT1/STAT2/IRF9 protein was incubated with GST-tagged Bclaf1 truncated proteins or GST control protein in PBS buffer with glutathione agarose (GE Healthcare) for 1 h at 4 °C. The incubated proteins were then washed and immunoblotted using anti-His or GST antibodies.

### Statistical analysis

Statistical analyses were performed using GraphPad Prism software to perform Student’s t test or analysis of variance (ANOVA) on at least three independent replicates. P values of < 0.05 were considered statistically significant for each test.

## Acknowledgements

We thank Drs. Huiqiang Lou (China Agricultural University), Jue Liu (Beijing Academy of Agriculture and Forestry Sciences), and Xiaojia Wang (China Agricultural University) for regents and cells. This work was supported by the National Key Research and Development Program of China (grant 2016YFD0500100), the National Natural Science Foundation of China (grant 31500703), and the State Key Laboratory of Agrobiotechnology (grant 2018SKLAB1-6).

## Author contributions

C.Q. and J.T. designed experiments; C.Q., R.Z., Y.L., A.S., A.X., M.W., W.H., and C.Y. performed experiments; W.F., and J.H. provided critical reagents and scientific insight; C.Q., R.Z., and J.T. analyzed data; C.Q. and J.T. wrote the manuscript.

